# Systems-Level Annotation of Metabolomics Data Reduces 25,000 Features to Fewer than 1,000 Unique Metabolites

**DOI:** 10.1101/155895

**Authors:** Nathaniel G. Mahieu, Gary J. Patti

## Abstract

When using liquid chromatography/mass spectrometry (LC/MS) to perform untargeted metabolomics, it is now routine to detect tens of thousands of features from biological samples. Poor understanding of the data, however, has complicated interpretation and masked the number of unique metabolites actually being measured in an experiment. Here we place an upper bound on the number of unique metabolites detected in *Escherichia coli* samples analyzed with one untargeted metabolomic method. We first group multiple features arising from the same analyte, which we call “degenerate features”, using a context-driven annotation approach. Surprisingly, this analysis revealed thousands of previously unreported degeneracies that reduced the number of unique analytes to ~2,961. We then applied an orthogonal approach to remove non-biological features from the data by using the ^13^C-based credentialing technology. This further reduced the number of unique analytes to less than 1,000.

## INTRODUCTION

It has become increasingly popular to perform untargeted metabolomics by using liquid chromatography/mass spectrometry (LC/MS). This is at least in part due to the large number of signals or features that are typically detected from most biological samples.^1–3^ While it is often assumed that these tens of thousands of detected signals provide “global” coverage of the metabolome, the number of metabolites being measured in an experiment has not been rigorously assessed. The major barrier preventing this type of analysis has been the challenge of identifying metabolites.^4^ To date, the overwhelming majority of the detected signals in any one untargeted metabolomics experiment have not been named. Even comprehensive efforts to identify as many metabolites as possible in a data set by using the most advanced informatic resources currently available have resulted in relatively small percentages of the total number of signals being identified.^5–7^ Thus, the basic question of how many unique metabolites are being profiled in an untargeted metabolomics experiment has remained outstanding.

It is important to note that uncertainties related to experimental coverage have not prevented the widespread application of the untargeted metabolomics technology. Improvements in instrumentation and software have made performing untargeted metabolomics with LC/MS relatively routine.^8^ Accordingly, the number of research cores offering LC/MS untargeted metabolomics services has increased dramatically over the last decade.^9^ The conventional workflows used by most research facilities, however, essentially sidestep the issue of experimental coverage.^10^ Their experimental output is a long list of signals or features, without thorough annotation. The data sets are either mined in a targeted fashion for specific metabolites with known retention times and fragmentation patterns, or only the small subset of signals that have a statistically significant difference between sample classes are further investigated.^11^ For many of these signals altered between sample classes, further investigation does not lead to identification because their accurate mass and fragmentation patterns do not match the accurate mass and fragmentation patterns of any known reference standard in metabolomic databases.^12^ Although it is common to refer to these unmatched signals as “unknown metabolites”, rarely is such a designation justified. Signals associated with contaminants, artifacts, and many adducts also do not return matches from metabolomic databases. These possibilities and others must be ruled out before gaining confidence that a signal is a bona fide, unique metabolite with an unknown structure.

The goal of the current study was to accurately enumerate contaminants, artifacts, and degeneracies at the systems level within the same data set to get an upper estimate of the number of unique metabolites detected in a representative LC/MS-based metabolomics experiment. For the purposes of this work, contaminant refers to a detected signal that does not originate from the biological sample being measured (e.g., solvent impurities and plastic leechables). Artifacts refer to features detected due to informatic error. As an example, artifacts can be caused by baseline fluctuations and poorly resolved components.^23,24^ Finally, degeneracy refers to multiple signals arising from a single analyte. There are many causes of degeneracy including: fragmentation, analyte adduction with various charge carriers (e.g., a proton, sodium, potassium, etc.), and the detection of naturally occurring isotopes (e.g., ^13^C, ^15^N, etc.). ^16,17,25^ A final, largely under-annotated source of degeneracy is the adduction of an analyte with other species present, including other analytes or the chemical background.

Although some degenerate relationships are well known and commonly annotated with the approaches described above, the prevalence of many degenerate relationships has not been previously estimated.^14^ Here we introduce and apply an approach that recovers relationships implied by the experimental data, rather than relying on a hypothetical predetermined list as is typically done in metabolomics. The approach allows for more comprehensive annotation, especially in the case of under-annotated adducts that may be specific to a single laboratory or experiment.

## RESULTS AND DISCUSSION

### Generating a representative untargeted metabolomic data set

In untargeted metabolomics, signals are often referred to as features, a convention we will follow here. A feature is a detected ion with a peak shape, unique *m/z,* and retention time. To estimate the number of unique analytes detected in a representative untargeted metabolomic data set, we set out to annotate three types of features: (i) degenerate features, (ii) contaminant features, and (iii) artifactual features. This work attempts to accurately enumerate each of these types of features simultaneously for the same data set. We annotated degenerate features by using mz.unity and a new contextual approach to find degeneracies implied by the data. We annotated contaminant features and artifactual features by using the credentialing approach.^26^ A requirement of the credentialing approach is uniform ^13^C-labeling. Given that there are convenient and well-established methods to culture *E. coli* on a uniformly labeled carbon source, we chose to focus our work on *E. coli*. Although mz.unity and credentialing were previously introduced for the targeted analysis of features, here we have developed these resources for systems-level studies so that they can be utilized simultaneously to assess analyte content in untargeted metabolomic data sets.

Metabolites from *E. coli* cells were extracted and analyzed with an LC/MS-based untargeted metabolomics platform, as detailed in Methods. In brief, metabolite extraction was achieved by using a combination of methanol, acetonitrile, and water. Extracted metabolites were separated with reversed-phase chromatography prior to being analyzed in positive polarity by a Q Exactive Plus mass spectrometer. These experimental methods (or variations thereof) are commonly applied in untargeted metabolomics.^27−29^ To process the resulting LC/MS data, we employed a custom informatic workflow (Figure 1). The workflow used an iterative, two-phase peak detection process. An in-house model-based feature detection algorithm was run on each of five individual replicates. Many of the resulting features are inconsistent between replicates due to subtle differences in the chromatograms from each file. It is common for some peaks to go undetected, or some peaks to be integrated differently between runs.^30^ These errors make further analysis challenging because a one-to-one feature grouping cannot be specified between replicates, and the established groups contain artificial variation in feature areas. To refine the features detected in the five replicates, we utilized the Warpgroup algorithm.^30^ Warpgroup considers all files in concert to identify “consensus features”, a set of feature integrations supported by all replicates. The result is a near one-to-one matching of features between samples (Figure 2A-B) and decreased variation introduced by informatic processing (Figure 2C-D). The Warpgroup refined feature detection is highly sensitive, allowing the recovery of features that, when processed in isolation, would be challenging to detect (Figure 2E). Here, we retained only features with a signal-to-noise ratio >5 and a coefficient of variation <0.5 after Warpgrouping. This resulted in 25,230 high-quality features in our representative data set. It is worth mentioning that analysis of our data set with the standard XCMS software resulted in the detection of more features compared to Warpgroup (see Supplementary Table 1). These data show that our informatic methods are not contributing to atypically high feature counts.

**Figure 1:**
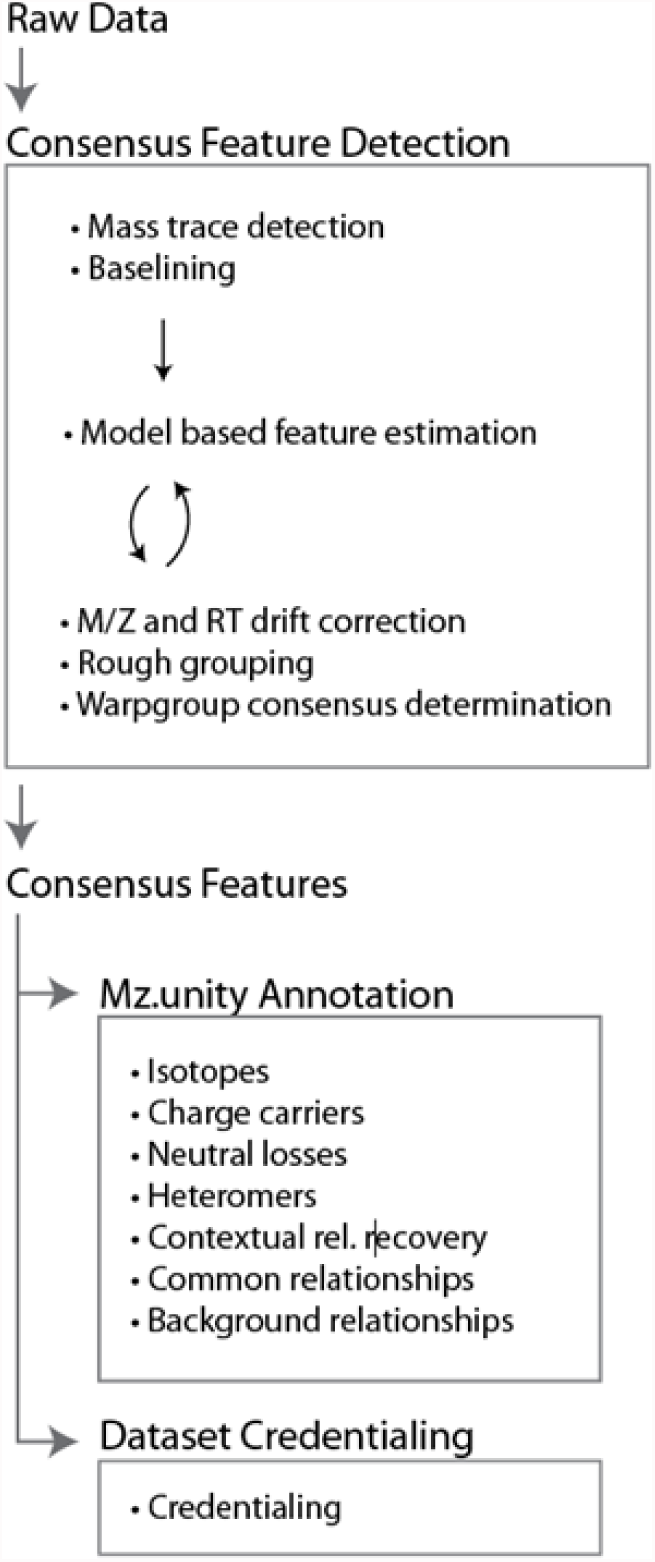
Our informatic workflow. Raw data were processed with in-house algorithms to first identify high-quality, consensus features (i.e., recurring features between replicates) and discriminate against processing artifacts. This consensus data set was further characterized by mz.unity (to estimate signal degeneracy) and credentialing (to estimate contaminants and artifacts).

**Figure 2:**
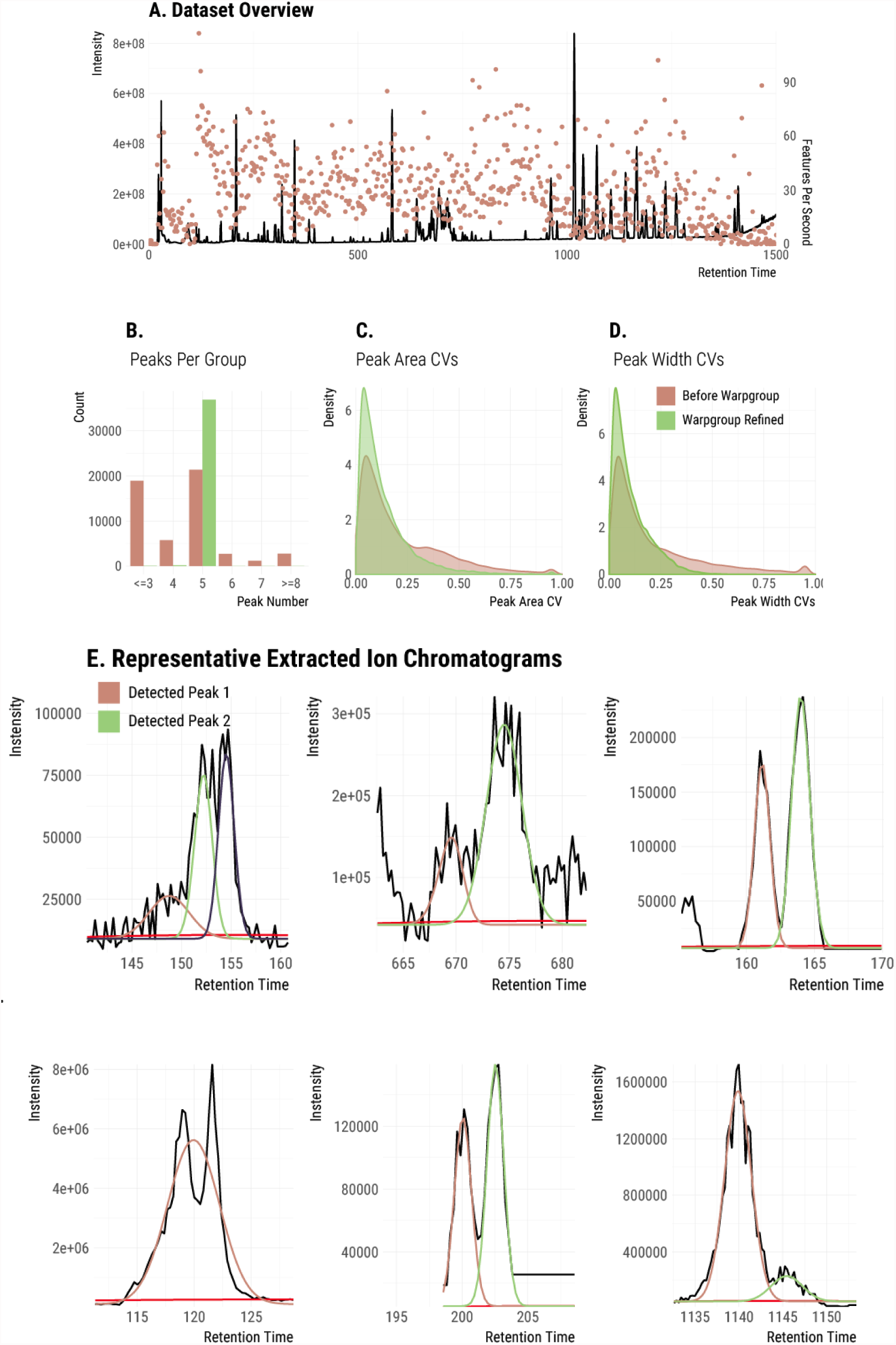
An overview of the consensus data set. (A) The base peak chromatogram of a representative run. The number of features detected during each second is overlaid. (B) The number of features detected in each group before (pink) and after (green) Warpgroup. Inconsistencies are resolved by Warpgroup. (C) The within group CVs of peak areas is decreased by Warpgroup. (D) The within group CVs of peak width are decreased by Warpgroup. (E) Several representative features detected by the informatic workflow. The estimated baseline is plotted in red.

We note that there is no universally accepted experimental platform for untargeted metabolomics at this time. The extraction techniques, chromatography, mass spectrometers, and peak detection algorithms used vary between laboratories and are often multiplexed.^31,32^ However, it is routine to detect tens of thousands of signals from a biological sample in most LC/MS experiments.^33,34^ Our detection of 25,230 consensus features from five replicates resulted in a data set with complexity that is typical of an untargeted metabolomics experiment.

### Simple annotations

As a first step to place an upper bound on the number of unique metabolites detected in our experiment, we performed a background subtraction. Specifically, we filtered features that were not at least two-fold higher than the signal detected in extraction blanks. These features represent contaminants or artifacts that are introduced during the sample extraction or data-processing steps. This reduced our list of 25,230 features to 12,797 (Figure 3A). Notably, this approach was developed to limit the number of artifact features returned by other processing software.

**Figure 3:**
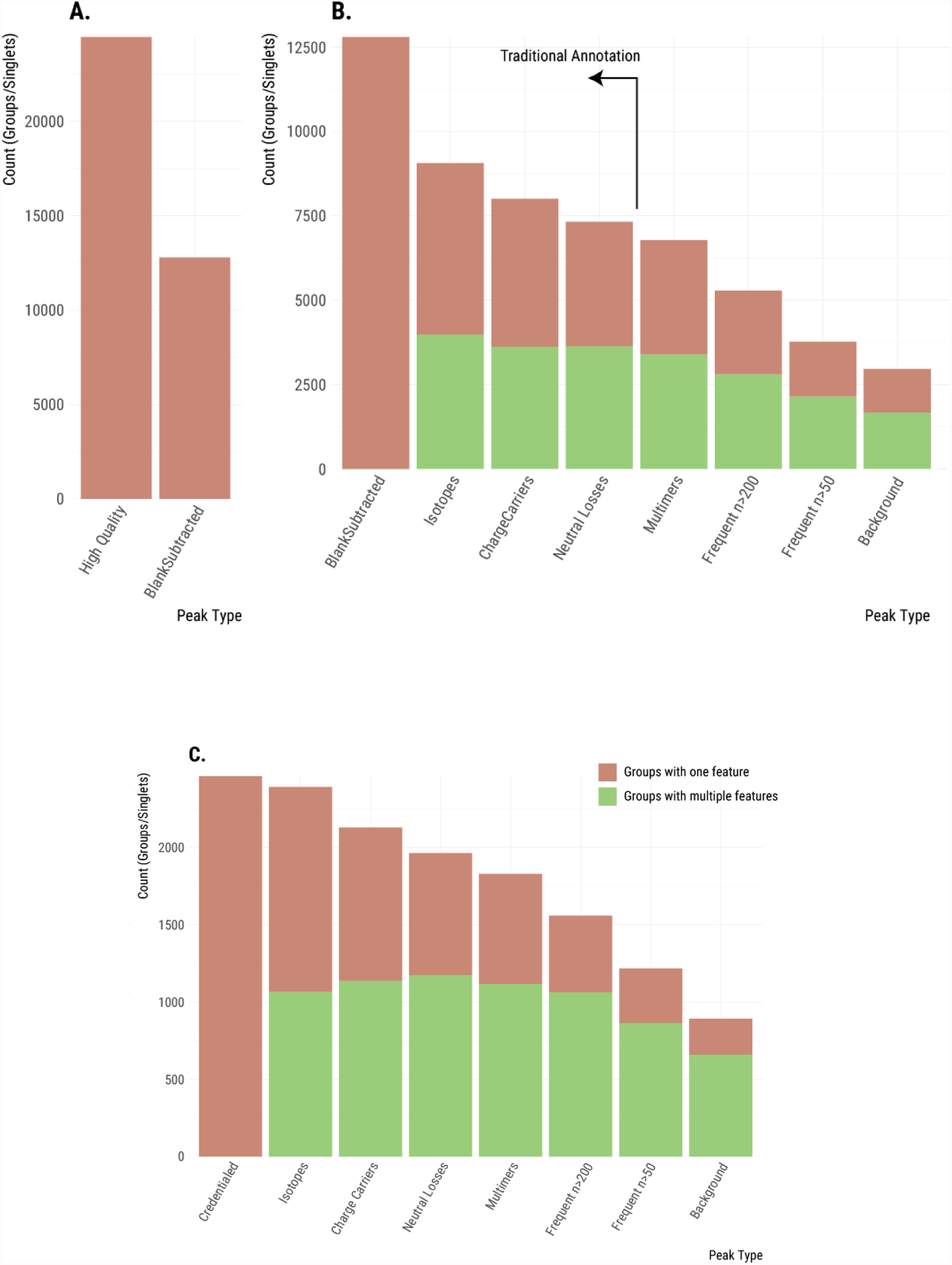
Plotting the maximum number of unique analytes detected throughout the steps of our annotation process. (A) Removal of features occurring in the blank. (B) Features are grouped as additional relationships are annotated. This reduces the maximum number of unique analytes. When a feature group contains multiple features, it is shown in green. When a feature group contains only a single feature (i.e., is a singlet), then it is shown in pink. Relationships from left to right: no relationships; isotopes; charge carriers; neutral losses; complex dimers (homo and hetero); frequent intrinsic relationships; situational adducts (background). (C) Similar annotation of features that were credentialed.

Next, we set out to annotate degenerate features (i.e., those features arising from the same analyte). We started our analysis by identifying simple relationships that are already commonly annotated in untargeted metabolomics.^14,35–37^ This included degeneracy due to carbon and other isotopes as well as common adducts and neutral losses. Annotations were made by using mz.unity, and degenerate features were grouped together.^17^ Because features within the same group arise from the same analyte, the number of “feature groups” provides a much better estimate of the maximum number of unique analytes detected in an experiment than the number of total features (Table 1 and Figure 3 B-C). In our subsequent descriptions, we will therefore transition from counting features to counting feature groups. A feature for which no degeneracy has been identified constitutes its own feature group, which we refer to as a singlet. Figure 3B shows the progressive decrease in the number of feature groups as isotopes, common charge carriers, and common neutral losses are annotated.

**Table 1:** A breakdown of the analyte number observed after each annotation step.

| Stage | Groups with more than one feature |  | Singlets |  |
| --- | --- | --- | --- | --- |
|  | All Features | Credentialed Features | All Features | Credentialed Features |
| Blank Subtracted | 0 | 0 | 12797 | 2462 |
| Isotopes | 3986 | 1066 | 5071 | 1326 |
| Charge Carriers | 3620 | 1137 | 4384 | 992 |
| Neutral Losses | 3640 | 1174 | 3678 | 790 |
| Multimers | 3400 | 1117 | 3381 | 712 |
| Commons n>200 | 2809 | 1063 | 2472 | 495 |
| Commons n>50 | 2149 | 864 | 1620 | 353 |
| Background | 1673 | 659 | 1288 | 233 |

When isotopes, common charge carriers, and neutral losses are annotated, the number of feature groups decreases from 12,797 to 7,318. We note that currently employed annotation approaches end here with the identification of simple relationships (see vertical line in Figure 3B). These results might suggest that there are as many as 7,318 unique analytes detected in the sample, but two observations suggested that much degeneracy still remained unannotated in our *E. coli* data set. First, about 50% of our feature groups still contain only a single feature (i.e., singlets with no detected relationships). Although in some cases singlets result from low-abundance analytes with no natural isotopes detected above noise level, the prevalence of singlets suggested that additional relationships remained unannotated. Second, we also know that the set of relationships annotated thus far are only a small subset of the possible degeneracies. A recent targeted study of glutamate demonstrated that many additional, complex sources of degeneracy can exist in LC/MS-based metabolomics that are not currently annotated with existing informatic resources.^17^ Glutamate was found to produce over 100 spectral peaks and exhibited complex adduct formation. Our objective was to comprehensively characterize these additional sources of degeneracy within a data set (*E. coli)* for the first time.

### Homo and hetero multimers

We then expanded our search for degenerate relationships to complex adducts (i.e., two or more species non-covalently bound to one another, such as dimers, trimers, etc.). Our search included analytes adducted with themselves (homo-relationships), as well as analytes adducted with different analytes (hetero-relationships). We considered all coeluting features as potential multimer partners evaluating all [m, z] values as possible adduct formers. The charge state was specified based on observed isotopes, or assumed to be a charge state of 1. As our conditions generally form ions with a single charge, we balance the +2 charge from the observed ions with the loss of a proton [1.00783, +1] for each multimer. Thus, a complex hetero-relationship between three detected features will satisfy: [m_1_, z_1_] + [m_2_, z_2_] – [1.00783, 1] = [m_3_, z_3_]. Grouping these detected complex adducts reduced the number of feature groups in our data set to 3,400 (see “multimers” bar in Figure 3B-C).

### Frequent intrinsic relationships show previously unannotated degeneracy

Current annotation approaches in untargeted metabolomics face the major challenge of determining the specific relationships to search for. While some relationships are well known and occur ubiquitously (such as the commonly annotated sodium or potassium adducts), constraining annotation to only these is significantly limiting. Other degenerate relationships are specific to experimental methodologies or the materials and reagents used during the analysis. Since there is no way to determine these relationships *a priori*, they have gone unannotated to date. Here we introduce an informatic approach to find data set wide, experimentally unique relationships that are implied by their context in the data. We then estimate their prevalence within our *E. coli* data set.

Common adducts and fragments will always coelute with the original analyte and will occur multiple times throughout the run.^17^ We leverage this fact and recover “frequent intrinsic relationships” by performing a frequency analysis of mass differences between all pairs of features eluting within one second of each other.^16^ Unrelated but coeluting analytes will exhibit mass spacing that is random and, as such, will not be enriched in the frequency distribution. Thus, frequently occurring mass differences represent probable degenerate relationships. Mass differences were calculated assuming a charge state of 1, a simplification that limits the analysis to relationships that do not include a charge-state conversion. A Gaussian kernel density estimation was performed on the observed mass differences with a bandwidth of 0.00001 Da (our observed scan-to-scan mass error) (Figure 4A). The heights of the local maxima represent the frequency and mass dispersion of each mass difference. Mass differences that are frequent and similar in mass will have large density estimates. The 24 most frequently observed mass differences are listed in Table 2.

**Table 2:** Recovered frequent intrinsic relationships. Not all recovered relationships shown were used in the annotation. The local maxima of the density are ordered by the number of occurrences. These frequently occurring differences are good candidates for peak relationships. Several well-known relationships are present, including alternative charge carriers at the top of the list.

| <u>$\Delta$ Mass</u> | <u>$\Delta$ Charge</u> | <u>Density</u> | <u>Known Species</u> |
| --- | --- | --- | --- |
| 21.9820 | 0 | 60.4 | $H^+ \leftrightarrow Na^+$ |
| 4.9554 | 0 | 55.2 | $NH_4^+ \leftrightarrow Na^+$ |
| 23.0760 | 0 | 33.6 |  |
| 18.0107 | 0 | 32.5 | $H_2O$ |
| 17.0266 | 0 | 30.5 | $NH_3$ |
| 28.0314 | 0 | 26.7 | $C_2H_4$ |
| 45.0580 | 0 | 23.4 | $C_2H_7N$ |
| 14.0157 | 0 | 23.2 | $CH_2$ |
| 65.1230 | 0 | 19.6 |  |
| 87.1046 | 0 | 18.2 | $C_5H_{13}N$ |
| 42.0470 | 0 | 16.6 | $C_3H_6$ |
| 44.0262 | 0 | 15.3 | $C_2H_4O$ |
| 39.9926 | 0 | 13.3 | $C_2O$ |
| 7.1020 | 0 | 13.1 |  |
| 15.9740 | 0 | 13.0 | $K^+ \leftrightarrow Na^+$ |
| 70.0783 | 0 | 12.5 |  |
| 29.0518 | 0 | 11.6 |  |
| 36.0713 | 0 | 11.3 |  |
| 15.9949 | 0 | 10.1 |  |
| 1.9967 | 0 | 9.3 | $^{41}K \leftrightarrow ^{39}K$ |
| 56.0627 | 0 | 9.3 |  |
| 12.9952 | 0 | 8.7 |  |
| 35.0373 | 0 | 8.7 |  |
| 20.9292 | 0 | 8.5 | $NH_4^+ \leftrightarrow K^+$ |

The effectiveness of the approach was confirmed by the recovery of two commonly known relationships as the most frequent relationships in our data set: the exchange of H^+^ and Na^+^ and the exchange of Na^+^ and NH_4_^+^. This result indicated that the analysis of frequent intrinsic relationships offers novel insight into the nature of features detected in metabolomic data sets. Notably, the approach returned a multitude of relationships that had not been included in our prior searches. These commonly occurring relationships are likely adducts or fragments, and may be specific to our sample or experimental equipment/materials. Figure 4B shows the peak pairs observed with mass difference [23.0760, 0] throughout the data set.

**Figure 4:**
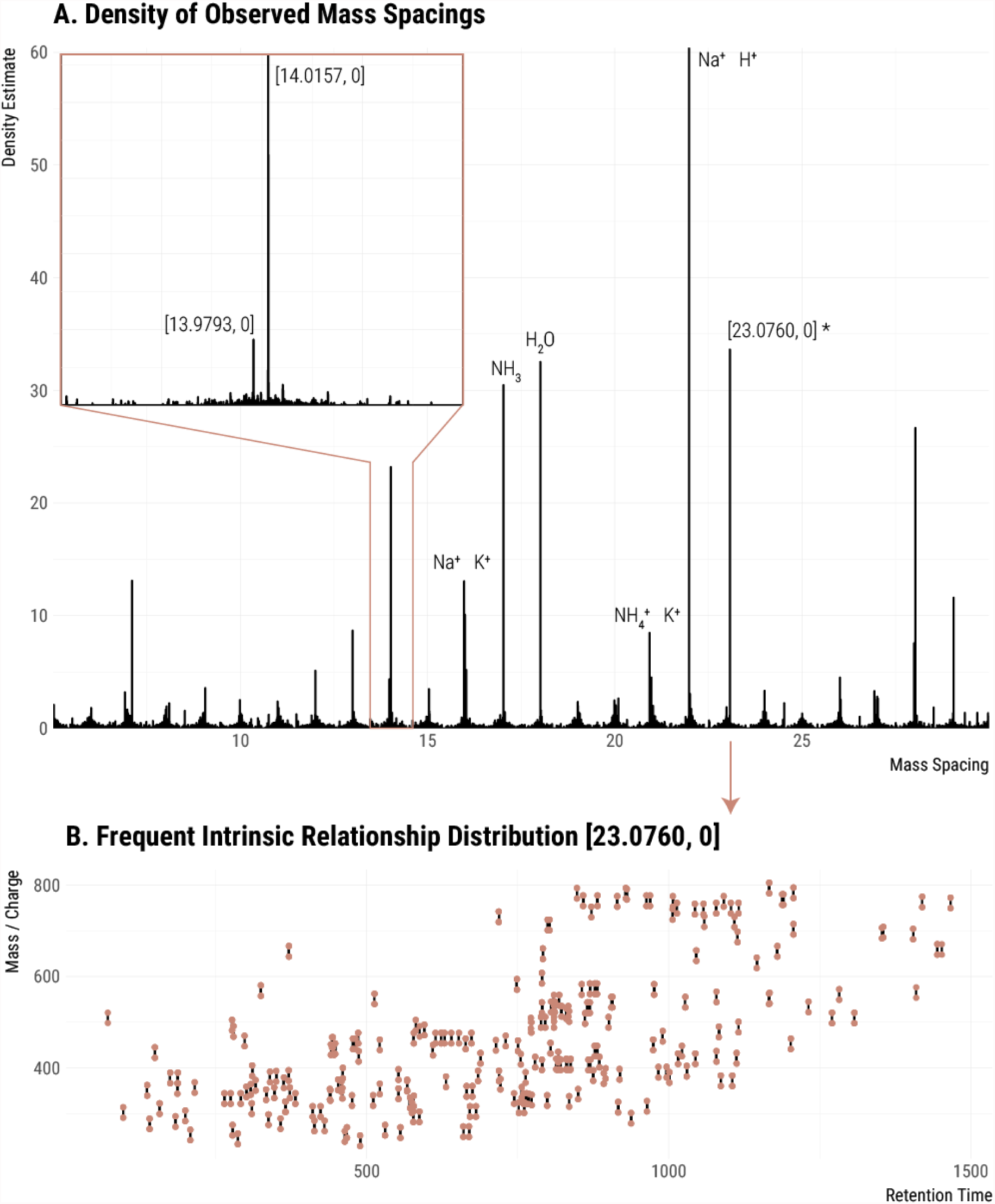
Detection of frequent intrinsic relationships. (A) The Gaussian kernel density of all pairwise peak relationships in the data set. Inset is a zoomed-in section around 14 Da. Known relationships are labeled with a formula. Unknown relationships are labeled with mass and charge transitions [m, z]. (B) Peak pairs of the recovered frequent intrinsic relationship [23.0760, 0] plotted in mass/charge and retention time (points). Line segments connect pairs with the specified spacing.

We recognize that the recovery of frequent intrinsic relationships can also return relationships between commonly coeluting, non-degenerate analyte pairs. Fully saturated and partially unsaturated lipids, for example, commonly coelute and have a mass difference of [2.0156, 0] (H_2_).^38^ We observed 176 occurrences of such a mass difference in our experiment. To minimize the risk of grouping unrelated features, we removed relationships with mass differences smaller than 15 Da and we applied two frequency cutoffs to illustrate the possible range of degeneracy. The conservative cutoff annotated and grouped frequent intrinsic relationships occurring more than 200 times (see bar labeled “commons n>200” in Figure 3B-C), while the aggressive cutoff annotated and grouped frequent intrinsic relationships occurring more than 50 times (see bar labeled “commons n>50” in Figure 3B-C). The inclusion of frequent intrinsic relationships in our data set annotation reduced the number of feature groups to 5,281 or 3,769, depending on the cutoff.

### Situational adducts due to background ions contribute significantly to degeneracy

To further expand the scope of our annotation, we considered a source of adduct ions that are present throughout the run: the chemical background. These ions lack a chromatographic peak shape, but they are detected throughout the experiment due to the ionization of solvents, their additives, or any contaminants present. Because the background ions coelute with every feature, it is reasonable to expect that they will produce many adducts. We refer to adducts between analytes and other presently observed species (such as background ions) as “situational adducts”.

A low-mass spectrum was collected, deisotoped, and background ions appearing at intensities higher than 200,000 were used as potential participants in situational adduct formation (Figure 5). Annotation of the identified situational adducts reduced our number of feature groups to approximately 3,000 (see bar labeled “background” in Figure 3B-C). This significant reduction in feature groups indicates that background ions are indeed a major source of feature inflation in our experiment. We also note that annotation of situational adducts reduced the number of feature groups containing only a single feature (i.e., singlets) to 1,288.

### Background ions give rise to some frequent intrinsic relationships

Some frequent intrinsic relationships that we detected are indicative of novel adduction or fragmentation phenomena in our untargeted metabolomic data set, and we were interested in the origin of these unknown relationships. We speculated that some of the frequent intrinsic relationships that we discovered were the result of analyte adduction with the chemical background described above. In the simplest of cases, we found that some frequently occurring mass-to-charge differences between features corresponded to the mass-to-charge values of background ions. However, we also noted that mass differences in the background ions were found in features. This indicated that a single analyte formed adducts with multiple background ions (Figure 5 and Figure 6) and therefore multiple situational adducts were detected for the same analyte. As the spacings between the background ions determine the spacings in the situational adduct features, we expect these repeatedly occurring spacings to be returned as frequent intrinsic relationships. Inspecting the returned frequent intrinsic relationships, we found several mass differences that also appear in the chemical background. This result is an additional confirmation of the effectiveness of frequent intrinsic relationship discovery and suggests that chemical background is a large source of feature inflation.

**Figure 5:**
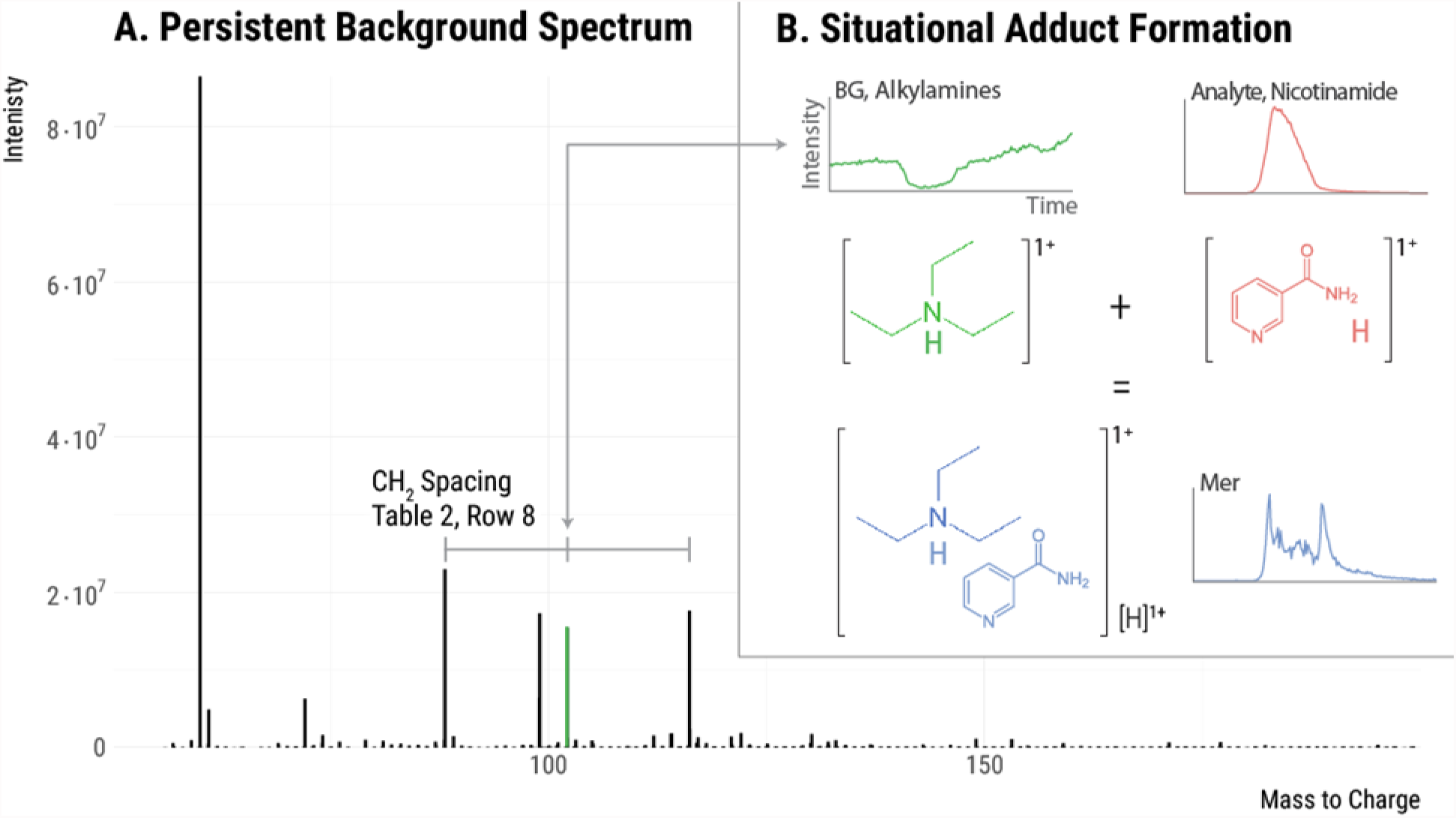
Situational adducts. (A) The persistent background spectrum observed in this experiment. The three indicated background peaks have mass spacings that correspond to a methylene group. These are likely an alkyl amine series with carbon numbers 5, 6, and 7. When these background species adduct with an analyte, situational adducts are formed. (B) An example of a situational adduct forming between background ion 102.1280 (a six carbon alkyl amine) and an eluting analyte. This process likely occurs with all three alkyl amine species throughout the run, giving rise to the frequent intrinsic relationships of mass 14.0157 (see Table 2, Row 8).

**Figure 6:**
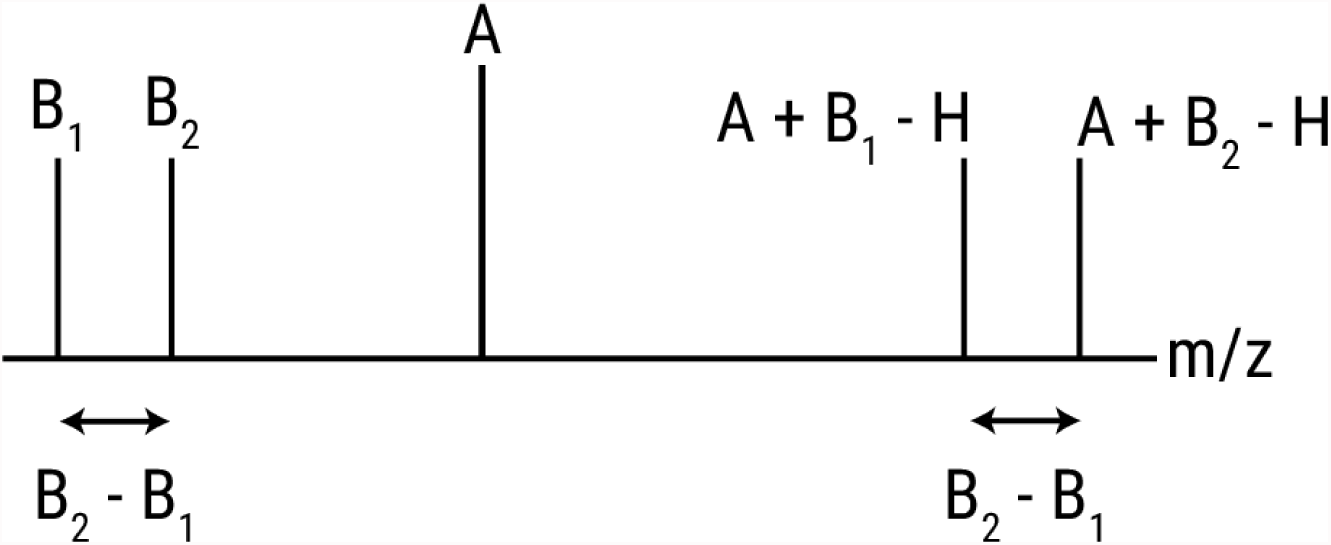
Schematic showing how background ions give rise to frequent intrinsic relationships. Analyte A is detected as an adduct of each background ion (B_1_ and B_2_). The spacing between the adducts (A+B_1_-H and A+B_2_-H) is equal to the spacing between the background ions.

We also performed formula decomposition on the frequent intrinsic relationships to further elucidate their origins. Interestingly, chemical formula CH_2_, C_2_H_4_, and C_3_H_6_ were found in the frequent intrinsic relationships exhibited by the chemical background. Additional analysis of the background ions indicated that they were an alkyl amine series. These alkyl amine species are known to form strong adducts and are commonly found as contaminants in alcohol solvents.^39^ We note that our laboratory has never performed ion-pairing experiments and the source of these reagents was solvent impurity as indicated by the series rather than sole presence of triethylamine. In developing our methods, we attempted to find solvents with the lowest possible levels of chemical background (Burdick & Jackson brand purchased from Honeywell). Unfortunately, alkyl amines seem to be ubiquitous in methanol and isopropanol LC/MS solvents.

### Removing artifacts and contaminants by credentialing

The degenerate relationships that we annotated above led to a striking reduction in the number of feature groups, indicating that fewer than 15% of the total 25,230 features that were detected in *E. coli* correspond to unique analytes. Even after this extensive annotation process, however, two sources of feature inflation remained in artifacts and contaminants. We applied an alternative experimental approach called credentialing to filter these features associated with artifacts and contaminants. The credentialing process introduces an isotopic signature into biological analytes during *E. coli* growth.^26^ Features in our data set displaying this isotopic signature are deemed “credentialed”, as they are known to be of *E. coli* origin. In contrast, features that do not display this isotopic signature are annotated as artifacts or contaminants. Credentialing does not rely on any of the relationship annotation approaches that we described above, and is thus an orthogonal and highly complementary approach to data analysis.

We first filtered non-credentialed features from the raw data set on the basis of isotopic signatures. The resulting set of features is free of artifacts, noise, and contaminants. This process returned 2,462 high-quality, credentialed features. We then took these credentialed features through the same annotation process as the full data set to remove degeneracy. Annotation of degeneracy reduced the estimated number of unique *E. coli* analytes being measured to 832 (Figure 3C).

## CONCLUSION

Detecting tens of thousands of LC/MS features from biological samples is typical in untargeted metabolomics, however, to date it has been unclear how many unique metabolites are actually being profiled. Our work here evaluated one representative untargeted metabolomics data set from *E. coli* to set an upper bound on the number of unique metabolites being measured. By using a new context-driven approach to identify degenerate features arising from the same metabolite, we determined that the ~25,000 features detected in our experiment corresponded to fewer than 2,961 unique analytes. An orthogonal and complimentary approach using credentialing isotope signatures to identify artifacts and contaminants similarly reduced the number of unique analytes detected. Out of the total ~25,000 features detected, only 832 passed both our degeneracy and credentialing filters.

We wish to emphasize that our work is unrelated to the size of the *E. coli* metabolome and should not be interpreted as an indication of the total number of intracellular metabolites present. There are certainly more than 832 *E. coli* metabolites.^44^ The purpose of our work is only to assess how many unique metabolites are being measured in a representative untargeted metabolomics experiment. Additionally, we note that our context-driven analysis of degeneracy is not exhaustive. Relationships that are uncommon and not indicated by background ions remain unannotated and may further reduce the number of unique analytes detected. Notwithstanding, our results suggest that there are an order of magnitude more features than unique metabolites in untargeted metabolomics experiments. This has important implications for designing untargeted metabolomics experiments and influences strategies for interpreting the data produced before establishing metabolite identifications.

## METHODS

### Materials

U-^13^C-D-glucose was purchased from Cambridge Isotope Laboratories Inc. (Andover, MA). *E. coli* strain K12, MG1655 was purchased from ATCC (Manassas, VA). Lennox LB broth powder and 5x M9 salts were purchased from Sigma-Aldrich (St. Louis, MO). Cell culture was performed with ultrapure water provided by a Milli-Q system (Millipore). LC/MS grade, Burdick & Jackson brand water, acetonitrile, methanol, and isopropanol were purchased from Honeywell (Morris Plains, NJ). Cortecs T3 reversed phase UPLC columns and column guards were purchased from Waters Corporation (Milford, MA).

### Generating credentialed samples

*E. coli* was grown in a rotary shaker at 37 °C and 300 rpm as previously described (Mahieu et al., 2014). A 100 mL volume of M9 minimal media was used with a glucose concentration of 2 g/L. Two cultures were grown in parallel, one using natural abundance glucose and a second using U-^13^C-glucose as the only carbon source. Cultures were grown to OD_600_ = 0.7, at which point they were harvested.

For harvest, flasks were removed from the shaker and placed on ice. The contents of each flask were pipetted into 50 mL conical tubes and centrifuged at 3200g and 4°C for 10 minutes. The supernatant was decanted and remaining media was gently rinsed off the top of the pellet with 0.5 mL of water. Conical tubes were then placed in liquid nitrogen and lyophilized for 24 hours, or until dry. This powdered, credentialed *E. coli* standard was then extracted to generate samples for untargeted metabolomic analysis.

Several replicate extractions were performed in parallel by using a previously described method.^26^ Briefly, five 2.5 mg samples of each ^12^C and ^13^C material were weighed out, while two empty tubes were included as extraction blanks. To these, 1,000 μL of 2:2:1 methanol:acetonitrile:water was added, followed by three freeze-thaw cycles with sonication and vortexing. After centrifugation, the supernatant was vacuum concentrated and reconstituted in 100 μL of 1:1 acetonitrile:water with internal standards. From these extracts, three samples were aliquoted for LC/MS analysis: natural abundance extract, a mix of 1:1 natural abundance extract and ^13^C extract, and the blank extract.

### Data set generation

Each sample was analyzed five times as analytical replicates. The untargeted LC/MS data set was generated in positive polarity on a Q Exactive Plus mass spectrometer with a HESI II source coupled to a Dionex 3000RSLC. The data set was collected with the following settings: aux gas, 5; sheath gas, 35; sweep gas, 2; capillary temperature, 300 °C; aux gas temperature, 200 °C; spray voltage, 3.5 kV; needle diameter, 34 ga; s-lens, 75 V; mass range, 100–1500 Da; resolution 70,000; micro scans, 1; max injection time; 100 ms; automatic gain control target: 1e6. Reversed-phase chromatography was performed with the Waters Cortecs T3 (2.1 mm x 50mm, 1.6um) column at a flow rate of 300 μL/min and a column temperature of 50 °C. Solvents were: A, water + 5mM ammonium acetate + 5uM ammonium phosphate; B, 9:1 isopropanol:methanol + 5mM ammonium acetate + 5um ammonium phosphate. An injection volume of 2 μL was used with a linear gradient of (minutes, %A): 0, 100; 28, 0; 30, 0; 30, 100; 35, 100.

Chromatographic features were detected by using a set of in-house algorithms. Mass traces were retained if they were longer than 10 scans, excluding missing peaks. Baselines for each mass trace were calculated by using the iterative restricted least squares method from the baseline R package. Model based peak detection was performed by using the skew normal distribution as a model peak distribution. This process resulted in a set of features detected in each replicate run. Features were grouped by mass and retention time using a density based method. Retention time drift and mass drift were corrected by fitting a loess curve of degree 2 to the distance from the mean value of each group against the mean retention time of each group.

Subtle variations from run to run cause many features to be integrated differently and sometimes not integrated in each file. Further, closely eluting peaks often lead to incorrectly grouped features. To resolve these missing values, refine the individual datasets and get a set of detected peaks consistent with all replicate runs, we applied the Warpgroup algorithm.^30^ Warpgroup is available at https://github.com/nathaniel-mahieu/warpgroup. Warpgroup takes as input the raw data and each file’s detected features combining them to output a set of consensus features. Parameters: sc.aligned.lim, 9; pct.pad, 0.1; min.peaks, 3. Of the detected peaks we retained only features with a signal-to-noise ratio >5 and a coefficient of variation <0.5 after Warpgrouping. This resulted in 25,230 “high-quality” features in our representative data set.

This consensus data set set is the standard output of an untargeted metabolomics experiment. As such, it was taken as a representative dataset for annotation of detected signals.

### Mz. unity based annotation

Mz.unity was applied to the dataset to detect mass and charge ([m, z]) relationships between eluting signals derived from a single analyte.^17^ We use [m, z] to denote the mass and charge of a species, where both are specified as opposed to *m/z* where the two are convolved. These searches find sets of features that have [m,z]s differing by a specific amount. Differences are specific to relationships, for example, loss of ^12^C and gain of ^13^C ([+1.003355, 0]), or loss of water ([-18.01057, 0]).

Searches were first performed for the following relationships: isotopes, common charge carriers, common neutral losses, and common adducts. We then searched for dimers between coeluting features. The dimer search posits each eluting [m, z] as a possible adduct former. The charge state was specified based on observed isotopes, or assumed to be a charge of 1. As dimers are normally formed with a charge from only one constituent, we also assumed the loss of a proton [1.00783, +1] for each pair.

Mz.unity is available at https://github.com/nathaniel-mahieu/mz.unity.

### Frequent intrinsic relationships

Groups of features eluting within 1 second of each other were taken, and their pairwise [m, z] differences were calculated after assuming a charge state of 1. A Gaussian kernel density estimation was performed on the mass differences with a bandwidth of 0.00001 Da (our observed scan-to-scan mass error). Local maxima of the density estimate were detected along with the estimated density at those locations. The heights of the local maxima represent the frequency and mass dispersion of each mass difference. Mass differences that are more frequent and more similar in mass will have larger density estimates.

We took enriched mass differences larger than 15 Da and occurring more than 50 times throughout the dataset into the mz.unity search.

### Situational adducts

Background ions that lack a chromatographic peak shape are an ever-present set of species that often form adducts with eluting analytes. These situational adducts are then detected as features having a chromatographic peak shape. A low mass background spectrum was collected, containing detected ions above 50 Da. This spectrum was deisotoped and background species appearing at higher than 200,000 intensity were used to seed possible adduct relationships. The [m, z]s of each background peak were included in the dimer search, as above after specifying the charge state based on observed isotopes or assuming a charge of 1.

### Credentialing

A high-confidence set of features were recovered from the ^12+13^C dataset by applying version 3.0 of the credentialing algorithm, which is available at https://github.com/pattilab/credential. Credentialing searches for pairs of peaks that have precise isotopic spacing expected from U-12C and U-^13^C analytes.^26^ This provides a filter against many forms of noise, contaminants, and artifact features. Credentialing was run with the parameters: ppmwid, 8; rtwid, 1.2; cd, 1.00335; mpc, c(12, 120); ratio, 1; ratio.lim, 0.1; maxnmer, 4. Credentialed features from the ^12+13^C data set were then matched to the ^12^C dataset by applying retention time and mass correction as above before grouping.

